# Dispersal overwhelms variation in host quality to shape nectar microbiome assembly

**DOI:** 10.1101/2023.01.05.522929

**Authors:** Jacob S. Francis, Tobias G. Mueller, Rachel L. Vannette

## Abstract

- Epiphytic microbes frequently impact plant phenotype and fitness, but effects depend on microbe community composition. Deterministic filtering by plant traits and dispersal-mediated processes can affect microbiome assembly yet their relative contribution is poorly understood.
- We tested the impact of host-plant filtering and dispersal limitation on nectar microbiome abundance and composition. We inoculated bacteria and yeast into 30 plants across 4 phenotypically distinct cultivars of *Epilobium canum*. We compared the growth of inoculated communities to openly visited flowers from a subset of the same plants.
- The abundance and composition of microbial communities differed among plant individuals and cultivars in both inoculated and open flowers. However, plants hosting the highest microbial abundance when inoculated did not have the highest abundances when openly visited. Rather microbial density among open flowers was correlated with pollen receipt, a proxy for animal visitation, suggesting a primary role of deterministic dispersal in floral microbiome assembly despite variation in host-quality.
- While host-quality can affect microbiome assembly, variation in dispersal was more important here. Host quality could drive microbial community assembly in plant tissues where species pools are large and dispersal is consistent, but dispersal may be more important when microbial dispersal is limited, or arrival order is important.

## Introduction

Phyllosphere microbes frequently influence plants’ expressed phenotype and ecological interactions. Plants benefit when microbes mitigate the effects of stress, enhance plant growth, or protect their host from antagonists (Stone *et al*., 2018). Yet other microbes are plant pathogens, deplete critical nutrients, or support the growth of other antagonists (Liu *et al*., 2020). Given the diverse and important effects of microbial communities on plant traits and fitness, understanding the processes driving plant microbial community assembly is a key goal. A predictive framework of plant microbiome assembly holds promise for both agricultural application (Busby *et al*., 2017; Toju *et al*., 2018) and deepening our understanding of ecological interactions in natural plant communities (Fitzpatrick *et al*., 2020).

Both deterministic and stochastic processes shape the assembly of plant microbiomes. Deterministic processes lead to predictable community trajectories given a set of ecological conditions (Vellend *et al*., 2014). In plants, deterministic microbiome assembly can be driven by variation in plant quality (Peiffer *et al*., 2013; Wagner *et al*., 2016; Leopold & Busby, 2020) that results in predictable variation in microbial survival, microbe-microbe interactions such as competition or priority effects (Fukami, 2015; Leopold & Busby, 2020; Mueller *et al*., 2022), or ecological interactions and their impact on microbiome (e.g. with herbivores Humphrey & Whiteman, 2020, or the environment Pusey & Curry, 2004; Gaube *et al*., 2021). Alternatively, stochastic or neutral processes that impact communities without regard to microbial species identity or host traits do not result in predictable community trajectories (Vellend *et al*., 2014). Ecological drift and stochastic dispersal generate non-deterministic variability in some microbiome communities (e.g. in *Arabadopsis*, Maignien *et al*., 2014; *C. elegans* Vega & Gore, 2017; and *D. melanogaster* Zapién-Campos *et al*., 2020). However, explicitly testing the relative role of deterministic and stochastic processes on plant microbiome assembly can be difficult (but see Edwards *et al*., 2018; rev. in Fitzpatrick *et al*., 2020). As such, the relative strength of deterministic vs stochastic processes in shaping plant microbiomes is still unclear (Dini-Andreote & Raaijmakers, 2018; Cordovez *et al*., 2019).

The floral microbiome has a central, unique, and often brief role in shaping plant fitness and ecology. Some pathogens use flowers to access plant vasculature (e.g. Anther-smuts and Erwinia Elmqvist *et al*., 1993; Sasu *et al*., 2010), sterilizing flowers or causing tissue death. Alternatively, non-pathogenic floral microbes are common (Vannette, 2020). In floral nectar, bacteria or yeasts may affect plant fitness via changes to floral phenotype that change pollinator visitation (Vannette *et al*., 2012; Schaeffer *et al*., 2017; Vannette & Fukami, 2018), shift pollinators’ on-flower behavior (Herrera *et al*., 2013; de Vega *et al*., 2022), or by competing with or facilitating other beneficial, commensal, and pathogenic microbes (Crowley-Gall *et al*., 2022; Mueller *et al*., 2022). Compared to the microbiomes of leaves or roots, floral microbiomes are highly variable among flowers on a plant, among individual plants, and among plant species (Rebolleda-Gómez *et al*., 2019; Vannette, 2020). Despite extensive documentation of this variability, whether deterministic processes are strong enough to overcome stochastic floral microbiome assembly or at what level of organization deterministic microbiome assembly varies (e.g. among individuals, among genotypes or cultivars, etc.) is not known.

Host selection is likely an important deterministic process in floral microbiome assembly (Rebolleda Gómez & Ashman, 2019) as floral microbiomes are not simply an unbiased subset of environmental microbes (Herrera *et al*., 2010; Rebolleda Gómez & Ashman, 2019; Rebolleda-Gómez *et al*., 2019). Individual plants can vary in their suitability for nectar microbes, for example apple cultivars differ in resistance to the florally transmitted pathogen *Erwinia amylovora* (Emeriewen *et al*., 2019). But less is known about filtering of commensal or beneficial nectar microbes and the mechanisms driving it. In lab experiments, floral traits can affect microbial survival and growth. Some nectar traits that could impact microbes are under genetic control, including nectar secretion rate, sugar concentration and composition, and amino acid concentration (Mitchell, 2004; Parachnowitsch *et al*., 2019; J Ryniewicz, M Skłodowski, M Chmur, A Bajguz, K Roguz, 2020). Additional nectar traits such as high sugar content (Herrera *et al*., 2010), the presence of secondary compounds and antimicrobial peptides (Adler, 2000; Palmer-Young *et al*., 2019; Christensen *et al*., 2021; Schmitt *et al*., 2021; Mueller *et al*., 2022), or biochemical conditions in nectar that generate reactive oxygen species (Carter & Thornburg, 2004) vary among plants and can impact nectar microbiome assembly. Because individual flowers are short-lived compared to other plant tissues, sometimes lasting only hours (Ashman & Schoen, 1994), small effects on microbial growth rate can have large implications for community assembly.

Dispersal is also a central mechanism in floral microbiome assembly that has both deterministic and stochastic components. In most floral communities less than half of flowers contain culturable yeasts or bacteria (Herrera *et al*., 2009; Vannette *et al*., 2021), which is generally attributed to dispersal limitation. Most nectar-inhabiting microbes depend on zoophilic dispersal (Vannette *et al*., 2021; except for some pathogens; e.g. Alexander, 1989), and clear vertical transmission of the nectar microbiome is not common. Dispersal can be deterministic if phenotypic differences among plants result in predictably different visitation by pollinators or other dispersers. There is some evidence for this: floral visitor networks predict the bacterial microbiomes of co-flowering plant species (Zemenick *et al*., 2021), and broad pollination guild can predict floral microbiome (de Vega *et al*., 2021). Nevertheless, plant-pollinator interactions are characterized by consistently low pollination driven by inadequate and partially stochastic pollinator visitation (Knight *et al*., 2005; Richards *et al*., 2009). Low dispersal probability may increase the relative importance of stochastic processes in microbiome assembly, and simulation modelling supports this hypothesis (Evans *et al*., 2017). Further, stochastic processes are more important for microbiome assembly in short-lived ecosystems like flowers (Zapién-Campos *et al*., 2020). Together, the short lifespan of flowers and stochasticity in pollinator visitation raise the possibility that dispersal-mediated impacts on microbiome assembly may be nearly neutral and might swamp out deterministic processes.

Here, we compared the relative strength of host plant selection and dispersal in shaping intraspecific variation in nectar microbiome. We experimentally tested for predictable variation in microbial growth and dispersal among individual plants and phenotypic cultivars. We inoculated a known microbial community into four phenotypically different cultivars of *Epilobium canum* where we restricted animal visitation. We examined if microbial growth and community composition was predicted by plant cultivar or individual identity when controlling for dispersal. We then compared the resulting inoculated communities with the floral microbiome of the same plants that were visited by pollinators. This allowed us to ask a series of 3 questions (Fig 1). First, does individual or cultivar-level variation in suitability for microbial growth have a deterministic impact on microbial community development (Study 1; Fig 1a)? Second, when we allow for natural variation in dispersal and host selection do plants or cultivars predictably differ in their microbial communities (Study 2; Fig 1b)? Finally, does the microbiome of openly visited flowers reflect microbial growth in inoculated flowers (Studies 1 and 2 combined; Fig 1c)? If floral filtering or host quality primarily shapes the nectar microbiome, we predict differences among individuals and/or cultivars in microbial communities independent of dispersal (in Study 1), and that these differences would reflect standing microbial communities in openly visited flowers (in Study 2). Alternatively, if deterministic dispersal is more important in shaping the nectar microbiome than floral filtering, we predict that plants or cultivars will have predictable microbiomes in openly visited flowers, but that the microbiome of open flowers would not reflect the microbiome of inoculated flowers. Finally, if stochastic processes were stronger than the deterministic portions of floral filtering or dispersal, we hypothesize that we could not predict the trajectory of microbial communities based on individual host or cultivar (i.e., no consistent individual or cultivar level variation in the microbiome in Study 1 and/or 2).

**Figure 1.**
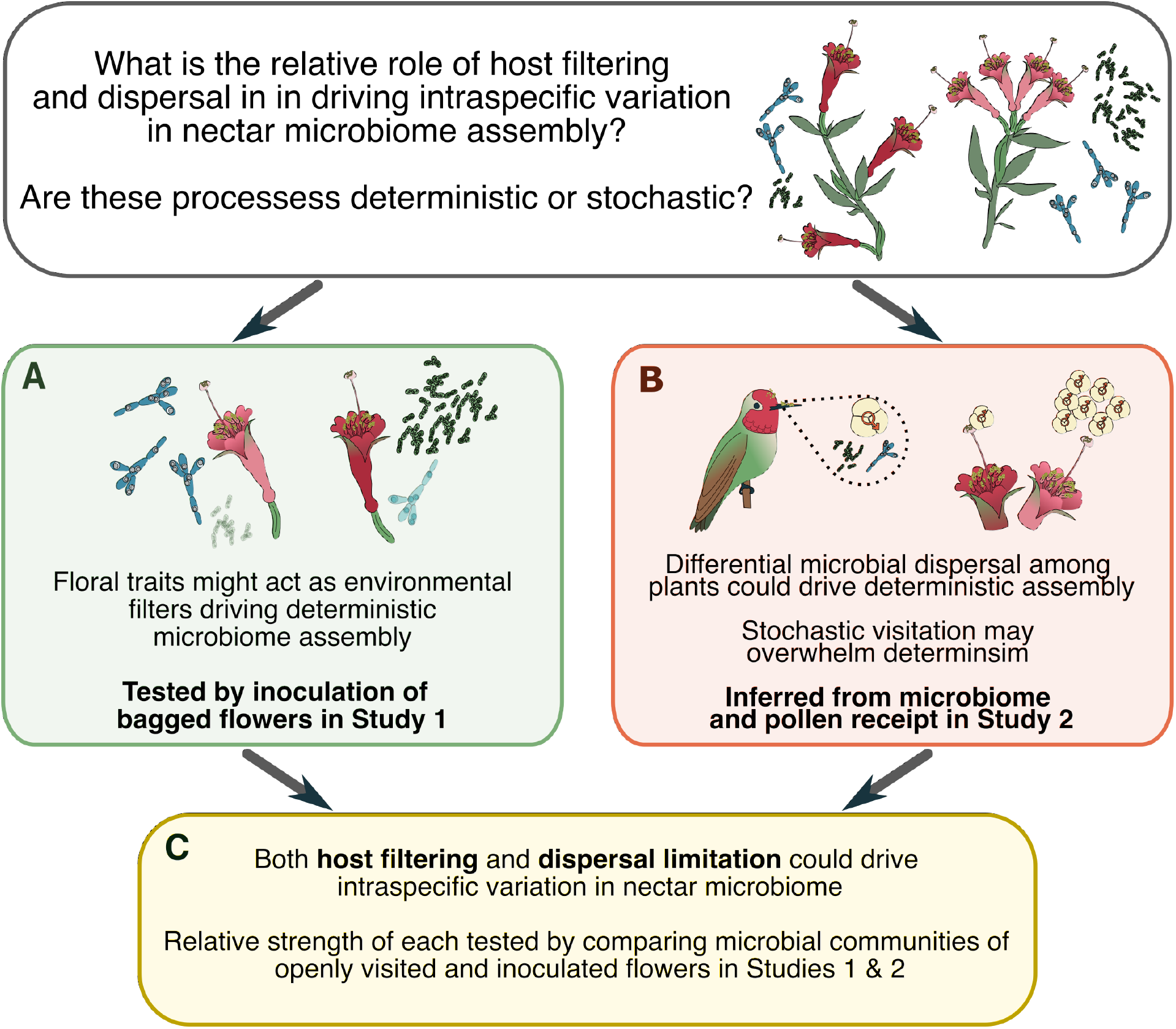
Conceptual diagram of hypotheses regarding the processes leading to intraspecific variation in nectar microbiome and how we tested them. First, we tested for cultivar and individual level variation in host filtering by inoculating a 2 species synthetic community into flowers where we restricted visitation (A: green box). Second, we tested for deterministic and stochastic differences among cultivars and in the standing microbiome of openly visited flowers (B: orange box). Finally, we tested whether plants that had high growth after inoculation also had high growth in openly visited flowers (C: yellow box).

## Materials and Methods

### Common garden design

To control for environmental influence on the nectar microbiome, variation in regional microbial species pools, or differences in the pollinator landscape, all plants were grown in a common garden on the campus of the University of California Davis (38.5371 N, 121.7728 W) embedded in a matrix of agricultural land. The garden consisted of 15 plots (7.6m x 4.6m), each planted with a community of co-flowering plants a year prior to the experiment (see S1 for a full species list by plot). At planting every bed contained 5 individuals of four morphologically distinct cultivars of the California endemic *Epilobium canum* (Greene, Onagraceae) including the wild accession *Epilobium canum* ssp. canum, (Canum); two horticultural cultivars: *E. canum* var Chaparral Silver (Silver, *E. canum), and* var Everett’s Choice (Everett); and the regional ecotype *E. canum* var. Calistoga (Calistoga). These four cultivars are phenotypically distinct and reflect either artificial or natural selection on standing phenotypic variation. Field work for experiments was conducted from 24 September 2020 – 26 October 2020.

### Study 1: Experimental inoculation

#### Inoculation Protocol

To test for differences among individual plants and cultivars we inoculated bagged flowers with a 2-species mixture of the yeast *Metschnikowia koreensis* and bacteria *Acinetobacter pollinis*, which commonly co-occur (Álvarez-Pérez & Herrera, 2013; Tsuji & Fukami, 2018). These strains were isolated from *E. canum* in the common garden and identified using MALDI-TOF using a custom built library of Sanger Sequenced microbial accessions (Morris *et al*., 2019 Bruker UltraFlextreme MALDI TOF/TOF). We created freezer suspensions (15% sucrose, 15% glycerol, 70% sterile ultrapure H_2_0) of this artificial microbial community made up of 5000 cells/µl of each species. Cells were quantified via hemocytometer. We created a single freezer stock at the beginning of the experiment, stored it at -80°C, and used aliquots across all inoculations to ensure that every flower was inoculated with the same initial microbial community. On the morning of each inoculation bout, we resuspended freezer stock in sterile 15% sucrose at 9:1 ratio giving us an inoculum with 500 cells/µl of each microbe. After thawing, the solution was vortexed for 30s. and stored for a maximum of 1-2h before inoculating flowers. We also created a control inoculum – the same sterile 15% sucrose used to dilute freezer stocks.

At least 48 hours before inoculation we removed all the male-phase flowers from a section of a plant and enclosed it in large pollinator exclusion bags (1 and/or 5-gallon paint filter bags, 200 microns, Cascade tools). We bagged flowers from 30 individual plants in Study 1. *Epilobium canum* is protandrous and takes approximately 2 days to proceed from male to female phase (Morris *et al*., 2019). By removing male flowers multiple days prior to inoculation, we could ensure that any male-phase flowers opened within our pollinator exclusion bags. Bags were effective at excluding large visitors to *Epilobium* (e.g., hummingbirds and bees), but less so for smaller animals (e.g., thrips and ants).

We randomly selected 14 male phase flowers on each plant for inoculation with microbial suspensions or control solutions (210 flowers of each treatment across the study). Unmanipulated bagged flowers contained on average 13.2 μL of nectar. Using sterile 10 µl microcapillary tubes (VWR, Radnor PA) we added 4µl of experimental solution to each flower (2000 cells each of *M. koreensis* and *A. pollinis*) or 4 µl of sterile 15% sucrose solution to controls, and flowers were marked using numbered jeweler’s tags. To estimate background microbial dispersal and contamination in bagged flowers, we sampled flowers inoculated with sterile control solutions (Supplemental Table S2). We inoculated flowers between 9:30 and 11:00 am across three days. After inoculation, all bags were replaced on the plants. During inoculation, we excluded any flowers that we observed being visited by animals while the bag was removed for experimental manipulation.

#### Assessing Microbial Community

We used culture-based methods to quantify microbes in inoculated flowers and control flowers as the two focal microbes form countable colonies on standard media (Morris et al. 2021). For inoculated and control flowers we excised flowers 72 hours after inoculation, transported flowers to the lab in coolers and extracted nectar in sterile condition. Not all flowers we inoculated persisted on the plant for 72 hours (N_control_=172 [Calistoga=44, Canum=43, Everetts=43, Silver=42] and N_inoculated_=173 [Calistoga=47, Canum=44, Everetts=43, Silver=39], ∼81% of inoculated flowers in each treatment persisted).

We collected nectar using 10µl microcapillary tubes, and measured nectar volume to the nearest 0.05 µl. If flowers contained more than 2µl of recoverable nectar, we destructively measured the sugar concentration on 1 µl of the sample (to the nearest 0.5% brix) using a handheld refractometer. The remaining nectar was diluted in 20 µl of sterile ultrapure water, diluted 10x further, and an aliquot plated on yeast media agar (containing 0.1 mg/ml chloramphenicol to control bacterial growth) and fructose-supplemented tryptic soy agar plates (containing 0.1 mg/ml cycloheximide to control fungal growth). Plates were incubated at 26°C for 48 hours and colony forming units (CFUs) counted. For the few plates (∼4%) which generated uncountable colonies, a single researcher determined the proportion of plate surface covered from which we estimated CFU values for each plate type (e.g. Tryptic Soy Agar or Yeast Media Agar). Finally we accounted for dilution to calculate CFU/μl for each nectar sample.

### Studies 2 and 3: Open flower sampling

We sampled 320 openly visited, non-bagged, female phase flowers across 7-8 individuals of each of the four cultivars above (10 flowers per plant, one individual Everett’s was unintentionally sampled twice, 8 days apart. This left us with 7 plants, one with 20 flowers). We sampled nectar and plated it to estimate microbial presence and abundance, as described above. We also removed stigmas of openly visited flowers immediately after removing flowers from plants and stored them in 70% ethanol to quantify pollen receipt.

### Studies 1 and 2: Measuring Floral Traits

We assessed nectar Brix of all samples that contained greater than 3 μl using a handheld refractometer. In 13 samples the measured Brix was greater than 50% sucrose. For these samples we tested the sugar concentration of the nectar sample mixed into 20ul of DI water (see inoculation data for samples). In openly visited flowers we also measured flower length (from the distal tip of the ovule to furthest distal petal tip) and width (the widest point between petal tips) of each flower.

### Assessing Pollen Receipt

To detect if flowers received pollen and to infer animal visitation, we used the stigmas of openly visited flowers to assess conspecific and heterospecific pollen receipt. We mounted each stigma in phenol-free fuchsin gel (Kearns & Inouye, 1993) and stored at 80° C. Stigmas were stored in 70% ethanol when flowers were collected in the field. We centrifuged the stigma-storage ethanol for each sample (1.5 min at 16000g), discarded the supernatant, resuspended the pellet in 70% ethanol, and mounted this solution in fuchsin gel to count any pollen grains that may have rinsed off stigmas in storage. *Epilobium* pollen is morphologically distinct from the pollen of co-flowering species, allowing for quantification of conspecific and heterospecific pollen receipt (Supplemental Fig 1). Although *Epilobium* flowers bear both male and female reproductive parts and produce copious amounts of pollen, flowers display spatial and temporal herkogamy (separation of anthers and stigma) which reduces within flower self-fertilization. However, we could not definitively identify whether conspecific pollen was self or outcrossed. While pollen receipt is not a perfect proxy for pollinator visitation, animal visitation to *E. canum* increases conspecific pollen deposition on stigmas and seed set (Snow, 1986).

### Statistical Analyses

All statistical analyses were completed in R 4.1.2 (R Core Team, 2014). Broadly, we used log-linear models (base R) and log-linear mixed effects models implemented in lme4 (Bates *et al*., 2014) to assess differences in microbial abundance among cultivars and plants. When testing for differences among cultivars we included a fixed effect of sampling date and a random intercept of plant to account for repeated measures on individuals. We included fixed effects of date to account for temporal differences in the pollinator community or any other unmeasured time-varying factors. We also ran separate models testing for differences in microbial communities among plots. If these tests indicated that spatial location (i.e. plot) impacted microbial communities we included plot in models testing for effects of interest. We tested for the significance of fixed effects using likelihood ratio tests for mixed effects models and F tests in linear models (implemented with the function drop1). Broadly, we began with fully specified models but dropped non-significant terms for reported statistical values.

### Study 1: Experimental inoculation

To examine host effects on microbial growth, we tested whether plant individual or cultivar explained variation in microbial growth (log CFU/µl of yeasts or bacteria +1, and total CFU/flower of yeasts or bacteria), accounting for repeated measures on individuals. To test whether *M. koreensis* and *A. pollinis* growth were correlated at the flower or plant level we regressed log CFU/µl *M. koreensis* +1 against log CFU/µl *A. pollinis* +1 in each flower against each other. To assess if inoculated yeast and bacteria respond similarly to variation in plant-level traits and test whether any plant was a universally good host, we regressed estimated marginal mean yeast and bacterial densities from the plant level analyses against each other. We used estimated marginal means to get predictions of yeast or bacterial growth after accounting for significant variation in covariates such as plot or sampling date. Plot was not predictive for bacterial densities, so we excluded it from subsequent models. Alternatively plots differed in yeast densities so we included it in individual models.

To examine if plant-level nectar sugar concentration predicted microbial growth, we first tested if cultivars and/or plants varied in Brix measured in uninoculated (control) flowers using a linear mixed effects model with a random intercept of plant to account for repeated measures in the cultivar model. We then tested whether estimated marginal mean Brix in control flowers from the plant level model predicted microbial community growth by regressing it against plant-level estimated marginal mean *M. koreensis* or *A. pollinis* densities. We only used control flowers that showed no microbial growth three days after inoculation (120 of 172 control flowers).

### *Study 2:* Open flower sampling

To examine if unbagged *E. canum* individuals varied in the probability of microbial colonization or microbial densities (log CFU/µl of yeasts or bacteria +1) we used two-stage hurdle models to account for zero inflation (45% open flowers did not contain microbes). We first used a binomial generalized linear model of the presence of culturable yeasts or bacteria across all sampled flowers. Subsequently only flowers containing culturable microbes were used to estimate log-linear models testing for differences between plants. To test for differences among cultivars we built similar models, but included a random intercept of plant to account for repeated measures on a single individual. To determine if plants and cultivars differed in pollen receipt we built similar 2-stage hurdle models.

To examine if flowers with pollen are more likely to contain microbes, we used chi-square tests comparing the presence of conspecific, heterospecific, or any pollen deposition with the presence of culturable yeasts, bacteria, or any microbes at the flower level (9 comparisons) and corrected for multiple comparisons using a false discovery rate. To test whether plants that received greater pollinator visitation also contain more flowers with microbes, we regressed the estimated marginal mean proportion of flowers that received conspecific or heterospecific pollen on a plant against the mean number of flowers containing culturable yeasts or bacteria. Finally, to test for pollinator-mediated selection on nectar concentration we regressed the mean nectar concentration in control plants against conspecific pollen receipt.

To test whether plant level traits including nectar sugar concentration (Brix) or volume, flower width or flower length impacted the probability of con- or hetero-specific pollen we used beta regressions (Grun *et al*., 2012), estimating separate models for each plant trait. Further we tested whether plant traits impacted microbial dispersal by building similar beta regressions of the above plant traits predicting the proportion of flowers on a plant that contained fungi or bacteria.

### Study 3: Does variation in inoculated microbial communities predict microbial density in open flowers?

To test whether plants that had the highest microbial densities after inoculation (Study 1) also had the highest microbial densities when openly visited (Study 2), we constructed 2 linear comparing the estimated marginal mean microbial densities in inoculated and open flowers at the plant level. We z-transformed and centered modeled densities around 0 to account for differences in total microbial densities in studies 1 and 2. Twenty-one plants were represented in both the inoculated and open flower data sets. By using the estimated marginal means from the density stage of the two-stage hurdle models in study 2, this model only includes flowers from study 2 that had successful microbial dispersal to them.

## Results

### Study 1) Do plants or cultivars predictably differ in microbial growth (independent of dispersal)?

Individual plants differed in their final densities of yeasts and bacteria 72 hours after inoculation with known microbial communities (Fig 2a,b). Of flowers inoculated, 98% contained culturable microbes (Supplemental Table 2). There was a 30-fold range among plants in the mean density of the yeast *M. koreensis* (log-linear model, F_29,141_=1.97, p=0.011, Fig 2a) and a 13-fold difference in the *A. pollinis* densities among plants (Fig 2b, log-linear model, F_29,138_=1.58, p=0.042). Generally, plant level differences were driven by a few plants that had significantly higher or lower microbial densities than others (Fig 2a,b). The results were qualitatively the same for total microbial cells per flower (See Supplemental Table 3 for models and p-values). *Epilobium* cultivars varied in final *M. koreensis* and *A. pollinis* densities (Fig 2a, log-linear GLMM, LRT, omnibus fungi χ^2^=8.21, p=0.042; bacteria LRT, χ^2^=9.68, p=0.021), but pairwise differences among cultivars were not strong (all post-hoc pairwise comparisons p>0.05). Plots also differed from each other in the density of *M. koreensis* (log-linear MEM, random intercept of plant, χ^2^=43.6, p < 0.001) but not *A. pollinis* (Table 1).

**Table 1:**
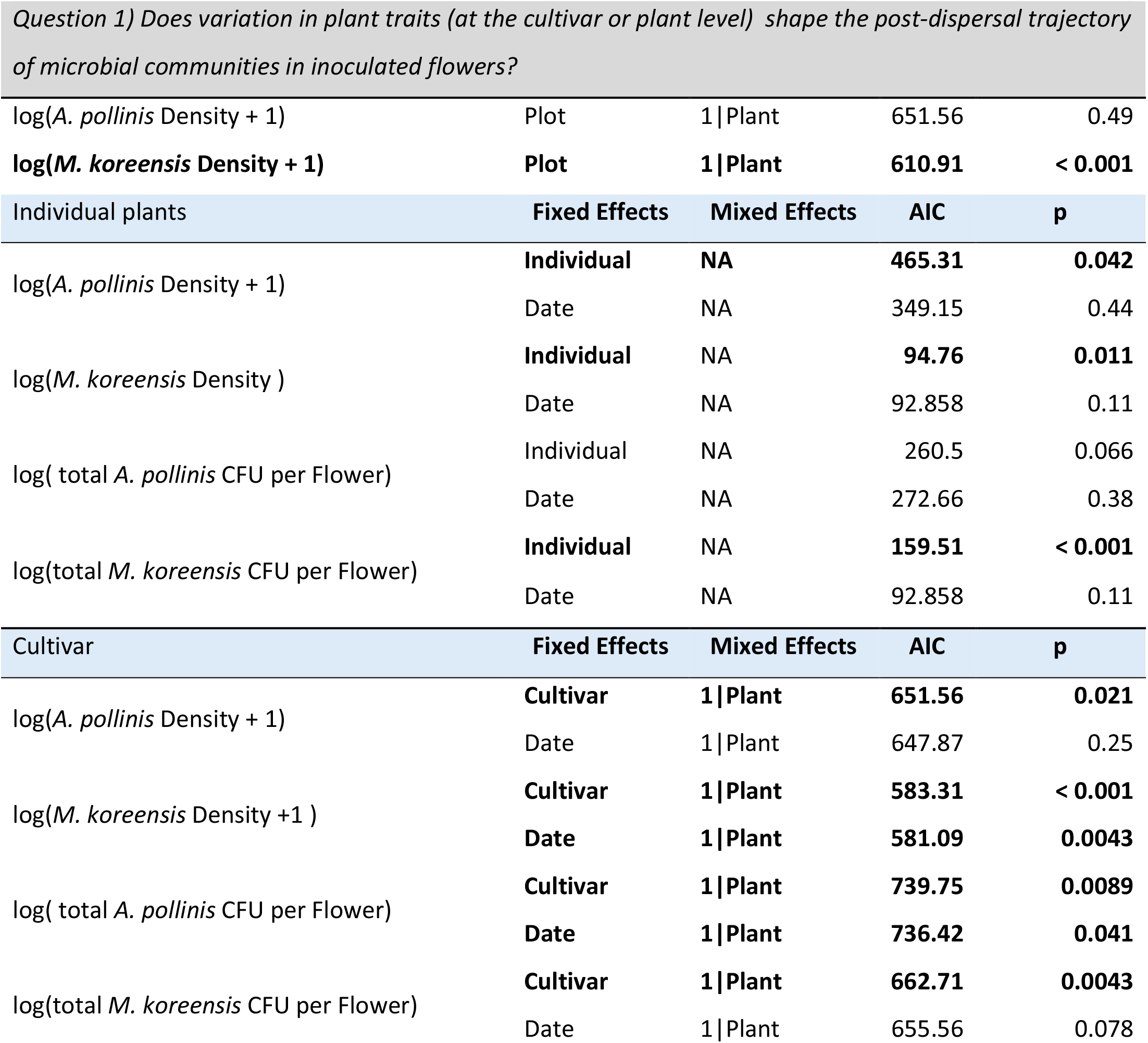
Models for inoculated flower experiment. Bold lines show significant fixed effects.

**Figure 2:**
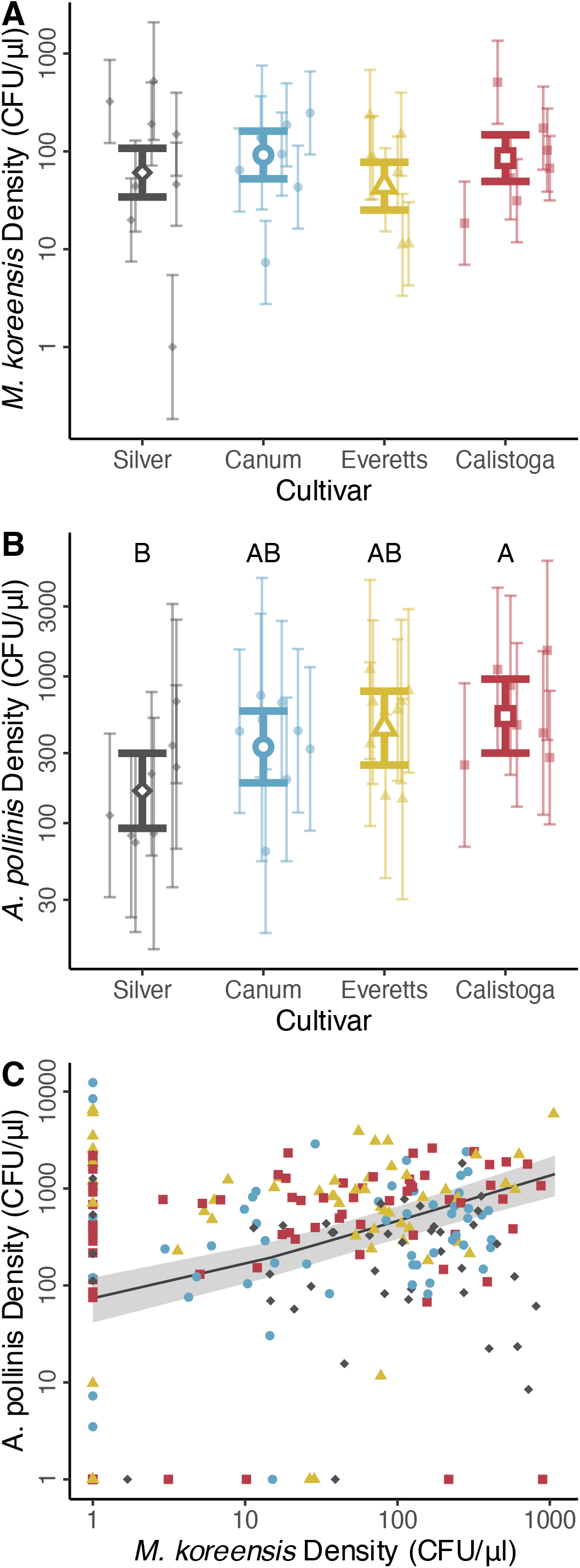
Density of (A) *M. koreensis* and (B) *A. pollinis* in inoculated flowers from Study 1. Small, closed points are the modeled mean +/- 95% confidence intervals CFU density for each plant (grouped, color and shape by cultivar). Large open points are the cultivar level modeled mean and 95% confidence intervals. There was significant variation among individual plants in both *M. Koreensis* and *A. pollinis*. Cultivars significantly differed from each other in *A. pollinis* densities (post-hoc significance indicated by letters). (C) Flower level correlation between *M. reukaufii* and *A. pollinis* with modeled relationship (black line) and 95% confidence intervals (grey fill). Points color and shape corresponds to cultivar. All densities shown on log scale. (Silver: grey diamonds, Canum: blue circles, Everett’s: yellow triangles, Calistoga-red squares).

Within a given inoculated flower, yeasts and bacteria densities were positively correlated, but the degree of correlation varied among cultivars (log-log linear mixed effects model with random intercept of plant, interaction term χ^2^=13.31, p=0.0040, fixed effect χ^2^=34.68, p < 0.001). In contrast, there was no significant correlation between *M. koreensis* and *A. pollinis* growth at the plant level (Supplemental Fig 2, log-log linear model, χ^2^=1.73, p=0.19).

We then examined if among-plant variation in nectar sugars was correlated with microbial growth. Individual plants differed in nectar sugars (Brix) in sterile control flowers (linear model, F_29,132_=3.74, p= p < 0.001; Fig 3a). This pattern was driven by a few individuals with very high or very low nectar sugar concentrations. Cultivars also differed in nectar sugar concentration (linear mixed model, χ^2^=14.57, LRT, p=0.0022 Fig 3a). Modeled mean plant-level sugar concentration did not predict *M. koreensis* (log-linear model, t=0.72, F=0.52, p=0.47, Fig 3c) or *A. pollinis* densities (log-linear model, t=-0.925, F=0.86, p=0.36, Fig 3b) at the plant level. Despite this, individual inoculated flowers with higher *Metschnikowia* densities also had greater nectar Brix (log-linear model, LRT, χ^2^=13.96, p < 0.001). This was not true for *Acinetobacter* (log-linear model, LRT, χ^2^=1.26, p=0.26).

**Figure 3:**
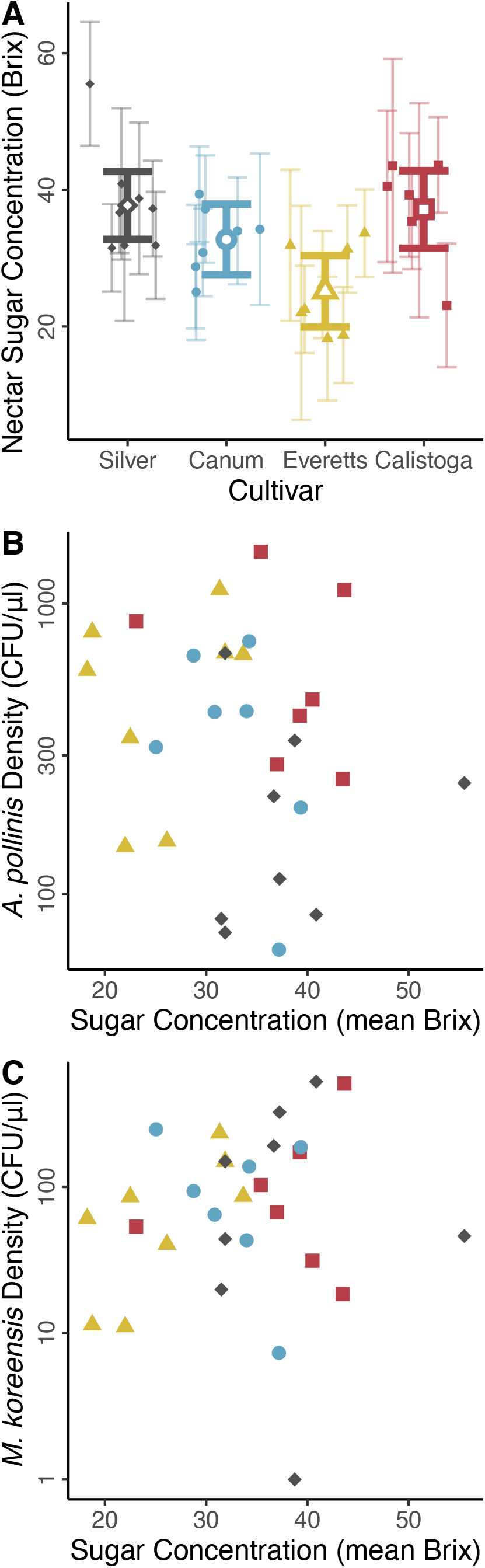
A) Nectar concentration (Brix) in control flowers varied among plants(small points) and cultivars (p = 0.0025, open points). Error bars represent 95% confidence intervals estimated from linear mixed effects model accounting for repeated measures on plants. There was no correlation between modeled mean *A. pollinis* (B) or *M. koreensis* (C) densities and modeled mean nectar concentrations at the plant level. (Silver: grey diamonds, Canum: blue circles, Everett’s: yellow triangles, Calistoga-red squares).

Sampling of uninoculated control flowers indicated that experimental inoculation was the main source of microbial inoculum in bagged flowers. Of uninoculated flowers, 10.0% contained fungi (mean 29 CFU/µl) and 25.9% contained bacteria (mean 712 CFU/µl) which we suspect may have been due to thrips visitation (Vannette *et al*., 2021). In contrast, 94.8%, and 94.7% of inoculated flowers contained fungi or bacteria with an average density of 363 CFU/µl and 1727 CFU/µl respectively (see Supplemental table 2 for breakdown of non-sterile controls by cultivar).

### Study 2) Do openly visited flowers predictably differ in microbial presence or growth?

Openly visited plants differed in their probability of containing detectable yeasts (binomial glm, LRT, χ^2^=69.99, p < 0.001) and bacteria (binomial glm, LRT, χ^2^=119.47, p < 0.001; Fig 4a,b). Additionally, cultivars differed in the likelihood that their flowers contained culturable yeasts (binomial glmm, LRT, χ^2^=14.37, p=0.002) and bacteria (binomial glmm, LRT, χ^2^=24.39, p= p < 0.001). Bacterial presence within cultivars varied among sampling dates and plots (binomial glmm, LRT, χ^2^=13.74, p=0.001 – date, and χ^2^=13.59, p=0.03 – plot), but not for yeasts.

**Figure 4.**
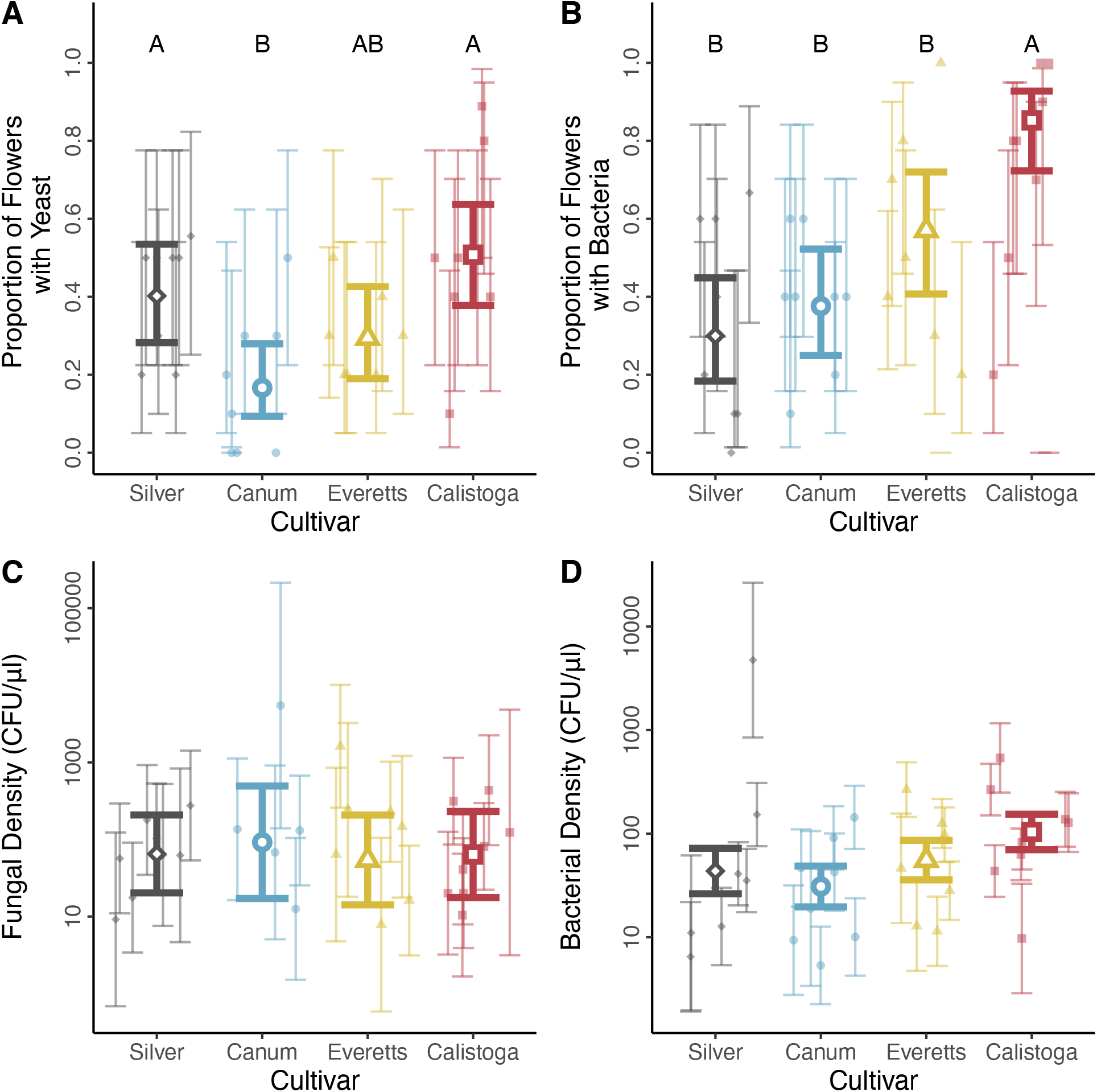
Modeled mean proportion of flowers that contain culturable fungi (A) and bacteria (B) by cultivar (large open points) and individual (small, closed points) colored and grouped by cultivar. Mean density of culturable fungi (C) and bacteria (D) in flowers that contained at least 1 CFU of each respectively by cultivar (large open points), and individuals (small, closed points) colored a grouped by cultivar. Error bars represent modeled 95% confidence intervals, note that density is log-scaled. (Silver: grey diamonds, Canum: blue circles, Everett’s: yellow triangles, Calistoga-red squares).

Among flowers that contained bacteria or fungi, plants differed in yeast and bacterial density (log-linear model, yeast F_27,82_=2.89, p=0.00017; bacteria F_30,126_=2.46, p=0.00032, Fig 4b,c). This was again driven by a few plants that had exceptionally high or low microbial densities. There were no significant differences among cultivars in non-zero yeasts or bacteria abundance (log-linear glmm, LRT, χ^2^=0.28, p=0.96; bacteria χ^2^=4.81, p=0.19, Fig 4b,c).

Next, we examined if plants or cultivars differed in their probability of pollen receipt and if pollen presence predicted microbial colonization at either the flower or plant level. Plants differed in the probability of receiving conspecific (binomial glm, LRT, χ^2^=111.32, p < 0.001) and heterospecific pollen (χ^2^=61.571, p < 0.001). Cultivars also differed in their probability of conspecific (binomial glmm, LRT, χ^2^=22.79, p < 0.001), and heterospecific pollen receipt (χ^2^=16.69, p < 0.001). There was temporal heterogeneity in heterospecific pollen receipt (χ^2^ = 13.77, p=0.0010). Flowers that received at least one conspecific pollen grain were more likely to contain bacteria (χ^2^=4.26, p=0.039), and to have microbe present at all (χ^2^=7.12, p=0.0076). However, all pollen receipt and microbe presence correlations became either non-significant or marginally significant when we corrected for false discovery rate (see Table 3). At the plant level however, the proportion of flowers that contained bacteria was positively correlated with the proportion of flowers that received heterospecific pollen (linear model, p=0.0037, t=2.82), but not conspecific pollen (p=0.74, t=0.30). The mean plant level pollen receipt (conspecific or heterospecific) did not predict the proportion of flowers that contained yeasts (Table 3)

**Table 2:** Models for open flower experiment. Bold lines show significant fixed effects.

| Question 2) Does intraspecific variation in plant traits or visitation shape the abundance and composition of microbial communities in openly visited flowers? |  |  |  |  |
| --- | --- | --- | --- | --- |
| Individual plants | Fixed Effects | Mixed Effects | AIC | p |
| Proportion of flowers containing bacteria | **Individual** | NA | **441.43** | **< 0.001** |
| binomial error structure | Date | NA | 381.96 | 0.063 |
| Proportion of flowers containing yeasts | **Individual** | NA | **410.14** | **< 0.001** |
| binomial error structure | Date | NA | 402.15 | 0.33 |
| log(bacteria per flower + 1) | **Individual** | NA | **209.11** | **< 0.001** |
|  | Date | NA | 197.2 | 0.71 |
| log(yeast CFU per Flower + 1) | **Individual** | **NA** | **161.93** | **< 0.001** |
|  | **Date** | **NA** | **158.26** | **< 0.001** |
| Cultivars | Fixed Effects | Mixed Effects | AIC | p |
| Proportion of flowers containing bacteria | **Cultivar** | **1 \| Plant** | **410.71** | **< 0.001** |
| binomial error structure | **Date** | **1 \| Plant** | **402.06** | **0.001** |
| Proportion of flowers containing yeasts | **Cultivar** | **1 \| Plant** | **405.25** | **0.0024** |
| binomial error structure | Date | 1 \| Plant | 400.29 | 0.55 |
| log(bacteria CFU per Flower + 1) | Cultivar | 1 \| Plant | 650.51 | 0.18 |
|  | Date | 1 \| Plant | 651.7 | 0.15 |
| log(yeast CFU per Flower + 1) | Cultivar | 1 \| Plant | 466.9 | 0.96 |
|  | **Date** | **1 \| Plant** | **480.82** | **0.0022** |

**Table 3:** χ^2^ tests of correlation between the presence of pollen and the presence of microbes at the floral level, and linear models at the plant level of probability of pollen receipt vs probability of microbe presence.

| Pollen Type | Microbe Type | p | Adjusted p |
| --- | --- | --- | --- |
| Conspecific | Microbes Present | **0.0076** | 0.069 |
| Conspecific | Bacteria Present | **0.039** | 0.17 |
| Conspecific | Yeasts Present | 0.11 | 0.35 |
| Heterospecific | Microbes Present | 0.44 | 0.54 |
| Heterospecific | Bacteria Present | 0.48 | 0.41 |
| Heterospecific | Yeasts Present | 0.2 | 0.54 |
| Any Species | Microbes Present | 0.33 | 0.49 |
| Any Species | Bacteria Present | 0.64 | 0.64 |
| Any Species | Yeasts Present | 0.23 | 0.41 |

| Plant Level |  |  |  |
| --- | --- | --- | --- |
| Pollen Type | Microbe Type | p | t-value |
| Conspecific | Bacteria Present | 0.74 | 0.30 |
| Conspecific | Yeast Present | 0.38 | 0.78 |
| **Heterospecific** | **Bacteria Present** | **0.0037** | **2.818** |
| Heterospecific | Yeast Present | 0.28 | -0.95 |

We did not find a strong signal of pollinator mediated selection on most plant traits measured (e.g. differences conspecific or heterospecific pollen receipt). But plants with longer flowers were more likely to receive conspecific pollen (beta-reg, z=5.00, p=0.01) (Supplemental Tab 3). In contrast, nectar and physical traits were associated with microbial incidence at the plant level. Plants with long and wide flowers were much more likely to contain yeasts (beta-reg, z=5.00, p < 0.001 and z=5.72, p < 0.001). Additionally, plants with higher nectar volumes had significantly higher incidence of bacteria (beta-reg, z=2.07, p=0.038).

### Study 3) Comparing between study 1 and study 2: Does microbial growth in inoculated flowers predict microbial presence or abundance in open flowers?

When comparing plant-level microbial densities between studies, there was no correlation between microbial growth in inoculated flowers and microbial density in openly visited flowers on the same plant. In other words, plant-level estimated marginal mean *M. koreensis* density in inoculated flowers was not associated with yeast density in open flowers (linear model, t=-1.15, p=0.27) nor was *A. pollinis* densities associated with mean bacterial densities in open flowers (linear model, t=-0.86, p=0.40).

## Discussion

Both dispersal limitation and floral filtering shaped nectar microbiome community assembly in this common garden of *Epilobium canum*, but dispersal dynamics masked differences in host-filtering under realistic pollinator visitation and microbial dispersal conditions. Despite strong host plant effects detected in study 1, the results presented here suggest that dispersal and stochastic processes (or unmeasured variables) are the primary factors affecting microbial presence and abundance in the *Epilobium* flower microbiome.

Host plant effects on the flower microbiome were significant in the absence of dispersal limitation. *Epilobium* plants differed in microbial communities when we controlled dispersal (Fig 2a,b), with plant identity explaining 25.5 and 52.9% of variation in *A. pollinis* and *M. koreensis* densities, and cultivar explained 6.72 and 38.7% respectively, suggesting that host genotype has a greater effect on yeast compared to bacterial growth in nectar. Genetically controlled host-filtering of plant microbiomes is well documented in other plant tissues (e.g. in leaves Bálint *et al*., 2013; and roots Xiong *et al*., 2021; rev. in Fitzpatrick *et al*., 2020). However host-mediated filtering of microbiomes may be lower or less detectable in flowers than in other tissues (Wei & Ashman, 2018) perhaps because some flower traits have high intra- and inter-individual plasticity in phenotype (e.g. nectar volume and concentration Nicolson & Thornburg, 2007) and nectar biomass is low. Despite clear deterministic plant level differences in microbiome abundance, the specific plant traits responsible are not clear. Nectar brix (a crude measurement of sugars) in uninoculated flowers did not predict microbial growth among plants for either microbe (Fig 3a,b). However, nectar is a complex mixture of mono- and disaccharides, free amino acids, and proteins (Nicolson & Thornburg, 2007) so future work should investigate how plant-level differences in nectar chemistry affect nectar microbiome assembly (Álvarez-Pérez *et al*., 2019; Mueller *et al*., 2022).

In addition to individual and cultivar level differences in microbial growth, we detected a strong positive correlation between *Metschnikowia* and *Acinetobacter* abundance in individual flowers (Fig 2c). Positive correlations between yeasts and bacteria within individual flowers have been detected previously, but they are not universal (Tsuji & Fukami, 2018; Álvarez-Pérez *et al*., 2019). A few hypotheses may explain this pattern. First, individual flowers on a plant may vary in quality, possibly due to variation in light, temperature, nectar secretion rates, or possibly even epigenetic mosaicism affecting floral traits (Herrera *et al*., 2021). Second, positive correlations may be due to co-dispersal, however, the pattern in the dataset reported here was in flowers where we controlled for dispersal. Finally, microbial metabolism may mediate facilitation within a flower, via the release of limiting nutrients (Christensen *et al*., 2021), detoxification of shared environments (Christensen *et al*., 2021; Mueller *et al*., 2022) or other mechanisms.

We saw strong evidence of deterministic dispersal limitation in microbiome assembly despite differences in host-suitability. Between 38 and 73% of open flowers on a plant contained bacteria and 17-50% contained fungi. This variation was not likely driven by host-filtering because 98% of inoculated flowers supported the growth of inoculated microbes. The incidence rates observed here reflect previous work surprisingly closely. In Mediterranean habitats in Spain, anywhere from 32-44% of flowers contain culturable yeasts (Herrera *et al*., 2009), and in the California coast range roughly 49% of flowers contain culturable bacteria and 20% contain culturable yeasts (Vannette *et al*., 2021). The results presented here suggest that intraspecific differences in microbial incidence can be as high or higher than differences among coflowering plant species. This intraspecific variation was not purely due to stochastic variation in dispersal; our phenotypically different cultivars differed predictably in realized microbial dispersal (Cultivar explained 22.1% and 9.74% of variance in bacterial and fungal incidence; Fig 4). Intriguingly, whether a plant had high incidence of bacteria or yeasts did not seem to be the simple product of high visitation by a single universal disperser because some cultivars had a high probability of yeast colonization but low probability of bacterial colonization (e.g. Silver, Fig 4). Given the significant but weak support for pollen receipt predicting microbial incidence these results suggest that different dispersal or establishment rates among microbes may be important (Vannette *et al*., 2021). Alternatively, these data suggest that flower visitors that disperse different microbial communities (i.e. hummingbirds, native bees, or honeybees Morris *et al*., 2019), also differ in pollination success. Moreover, the interplay between pollinator visitation frequency, nectar consumption and secretion rates, and microbial growth dynamics among flowers may contribute additional complexity beyond simple visitation dynamics.

Despite a weak signal of host filtering in openly visited flowers (Fig 4c,d), there was no correlation between the mean plant-level growth of *Metschnikowia* or *Acinetobacter* in inoculated flowers and the mean densities of culturable yeasts or bacteria in openly visited flowers on those same plants (whether we excluded non-colonized flowers or not Fig 5). At the individual level, plant identity was a strong predictor of the presence or absence of yeasts (R^2^ = 19.7%) and bacteria (R^2^ = 41.9%) and of fungal (R^2^ = 41.5%) or bacterial (R^2^ = 36.0%) growth in flowers that did have microbes present. One explanation for this pattern is that dispersal is a major driver of microbial community assembly in this system, overwhelming signatures of floral filtering. While the preeminence of dispersal in shaping floral microbial communities has been recognized in other plant species (Rebolleda Gómez & Ashman, 2019), we found strong evidence that it could drive intraspecific variation in microbiome assembly even at small spatial and short temporal scales.

**Figure 5:**
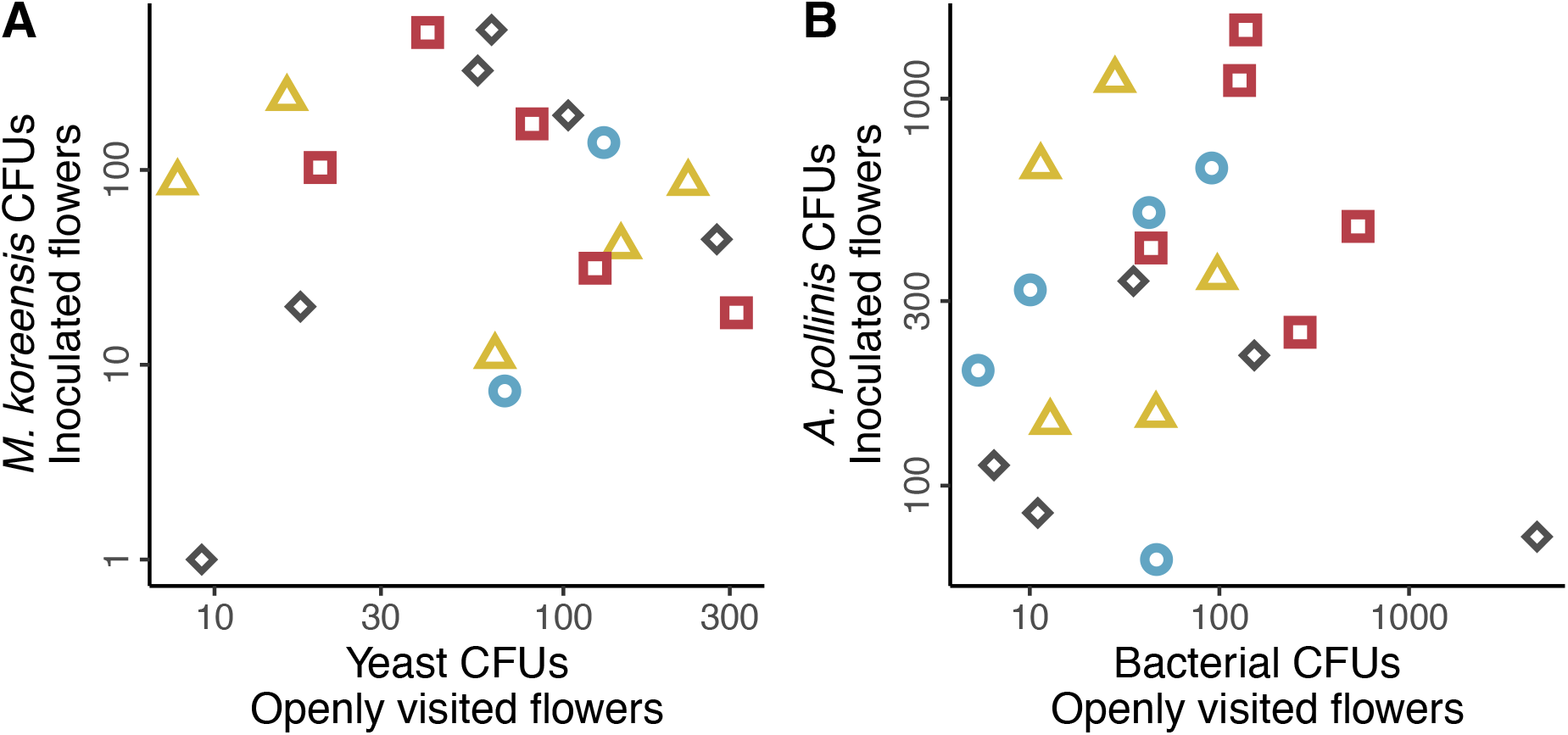
(A) There was no correlation between mean fungal CFUs in open flowers that had fungal cells present and *M. koreensis* growth in inoculated flowers on the same plant (p=0.20). (B) Similarly, mean bacterial CFUs in flowers that contained bacteria did not predict *A. pollinis* growth on the same plant (p=0.36). (Silver: grey diamonds, Canum: blue circles, Everett’s: yellow triangles, Calistoga-red squares)

We note that there are likely additional explanations for the incongruence in microbial abundance between studies one and two. One difference is that microbial communities in open flowers were likely more diverse than the inoculated two-species communities, and plant traits may vary in their effects on microbial species (Vannette & Fukami, 2018; Mueller *et al*., 2022). Further, in-flower interactions such as competition, facilitation, and priority effects can drive nectar community assembly (Fukami, 2015; Álvarez-Pérez *et al*., 2019b) and may be more pronounced when species diversity is higher. However, the *A. pollinis* and *M. koreensis* strains we selected are the most common morphospecies in our open samples. These species often co-occur (Álvarez-Pérez & Herrera, 2013) and are common and dominant in *Epilobium canum* (Morris *et al*., 2019), so our simulated communities are likely good, but simplified, representations of natural communities. Nevertheless, the complex multispecies interactions and historical contingencies acting in natural communities were absent in our inoculated flowers. Another difference between the studies is that the time since microbial dispersal also may have swamped out possible floral filtering in open flowers. Future work should use more complex microbial communities in inoculation trials to test if flowers selectively inhibit the growth of specific species or strains of microbes.

If dispersal dynamics swamp out host selection on microbiome assembly, can microbes mediate the evolution of floral traits? Some have argued that the microbiome is part of a flower’s extended phenotype and may impact floral evolution (Rebolleda-Gómez *et al*., 2019), but dispersal dependence in these systems makes predicting this evolution difficult. This extended phenotype is most likely to impact floral evolution if heritability of the microbiome is high. These results show that the floral microbiome may be heritable (e.g. deterministically dependent on plant traits) but likely has lower heritability than other genetically controlled floral traits which are not contingent on stochastic dispersal. For example, in our open flower data set, plant identity explained 56% percent of variation in control flower nectar concentration but only 36% and 42% of variation in bacterial and yeast densities and 41% and 20% of the variation in the probability that a flower contained yeasts or bacteria respectively. Because of the association between flower visitation and microbial presence, we predict that floral traits that impact pollinator attraction may be important in shaping the “heritability” of the nectar-microbial phenotype. We found some evidence for this here: plant-level sugar concentrations were positively correlated with the proportion of flowers containing bacteria. We also found that the presence of bacteria was weakly correlated with pollen receipt. Further the correlations between visitation and microbiome suggest a novel hypothesis: that plant species that have a high probability of adequate pollination in one or very few visits (e.g., plants with pollinaria or extraordinarily high Pollen:Ovule ratios), may be under less floral-microbe mediated selection because any microbes dispersed with pollen would not be able to impact fitness via changes floral phenotype that impact pollinator behavior. We predict that for microbes to act as heritable parts of a plant phenotype or shape plant trait evolution, microbial dispersal would have to be either consistently high (via microbe dispersal traits) or extremely costly/beneficial (as is the case with pathogenic microbes where the eco-evolutionary dynamic may be different Alexander, 1989; Elmqvist *et al*., 1993).

Taken together, our results suggest that floral microbiome assembly is contingent on both host selection and dispersal limitation and that the realized floral microbiome is a process of interactions between them and their relative strengths. Both deterministic and stochastic processes played a role in floral microbe community assembly here, and each may have differing importance among microbes and across scales (plant vs. population). Because floral microbes are dispersed primarily by animals who make predictable decisions based on plant traits, nectar microbiomes may be unique from other plant tissues because host selection can act not only on growth rates but also on deterministic dispersal probabilities. However, we suggest that the role of microbial dispersal limitation may be an underrecognized driver across other plant tissues. Recent evidence from phyllosphere microbes suggests that possible dispersal from co-occurring plants is an important factor in driving leaf microbiome assembly (Meyer *et al*., 2022). Further, new work suggests that at large biogeographic scales, rhizosphere bacteria and fungi can be dispersal limited, showing spatial autocorrelation despite microbes having broad ecological niches (Zhang *et al*., 2021). Our work adds experimental evidence that deterministic dispersal can overwhelm host selection in some cases (Cordovez *et al*., 2019) and that additional studies of plant microbiomes should consider this possibility.

## Acknowledgements

We thank the Vannette Lab (Danielle Rutkowski, Shawn Christensen, Amber Crowley-Gall, Marshall McMunn, Jake Cecala, and Dino Sbardellati) and Insect Ecology group (Sharon Lawler, Richard Karban, Emily Meineke, Neal Williams, and Jay Rosenheim, and members of their labs) who gave useful feedback on early manuscript drafts. David Fujino, Haven Kiers, Louie Yang, and Miles Daprato were integral in establishing the common garden used in this project. We acknowledge funding from NSF DEB 1846266 to RLV, NSF DBI 2109460 to JSF, and the Saratoga Horticultural Research Endowment to RLV. MALDI Biotyper funded by NIH S10 grant to UC Davis # S10OD018913-01A1.

## Author Contributions

All authors contributed to the conceptualization, data collection, and preparation of the manuscript. JSF performed statistical analyses with feedback from TGM and RLV, curated the data, and prepared the original draft of the manuscript.

## Data Availability

All data will be submitted to a public repository upon manuscript publication.

## Supplemental material

**Supplemental Table S1:**
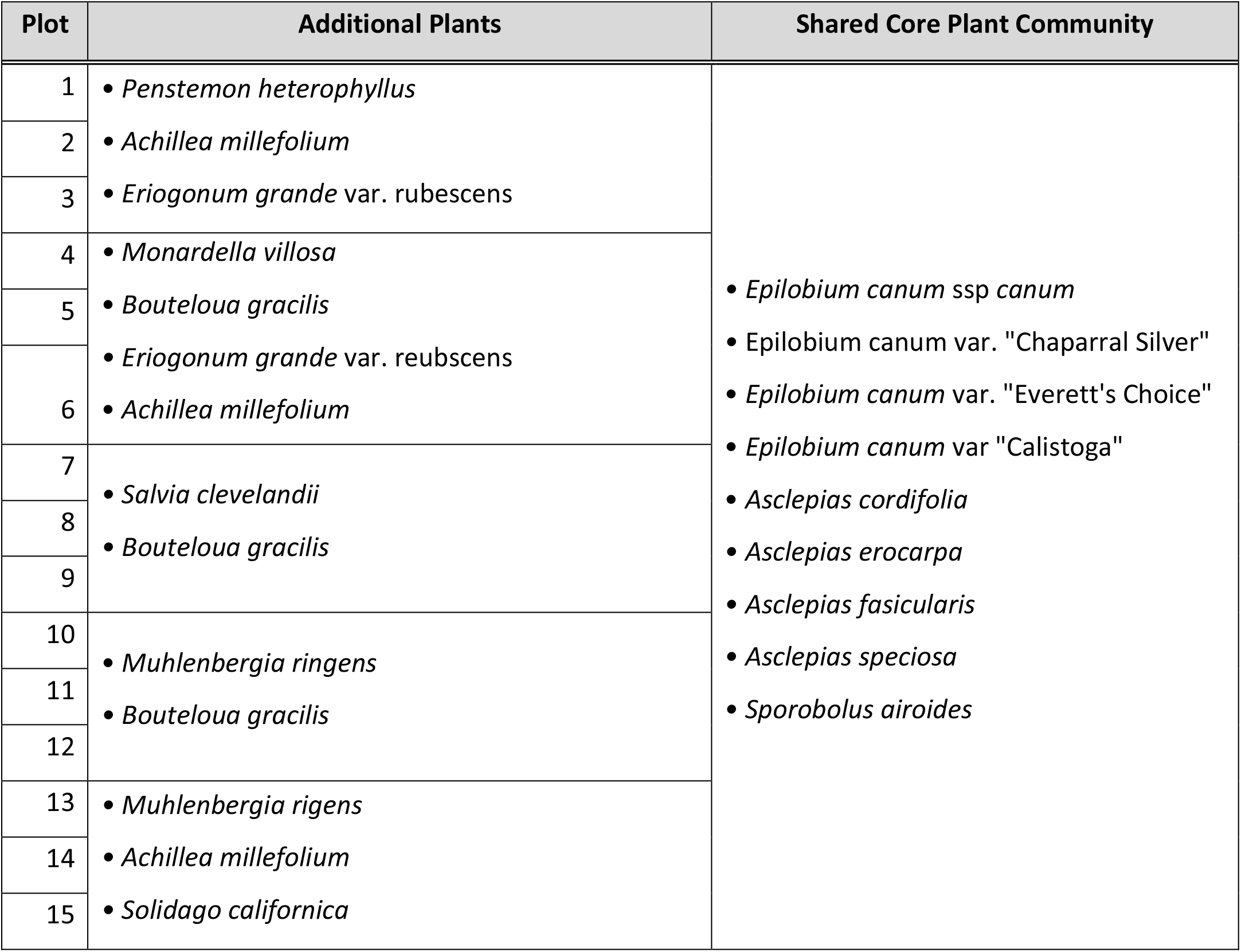
Plant community composition of each common garden plot at UC Davis. Each plot received the same watering regime.

**Supplemental table S2:** Microbial presence in control flowers from Experiment 1.

| Cultivar | Fungi<br>Absent | Fungi<br>Present | Proportion<br>Colonized (F) | Bacteria<br>Absent | Bacteria<br>Present | Proportion<br>Colonized (B) |
| --- | --- | --- | --- | --- | --- | --- |
| Calistoga | 36 | 8 | 0.26 | 24 | 20 | 0.46 |
| Canum | 39 | 4 | 0.11 | 34 | 9 | 0.21 |
| Everett | 39 | 4 | 0.11 | 33 | 10 | 0.23 |
| Silver | 40 | 2 | 0.05 | 37 | 5 | 0.12 |
| Total | 154 | 18 | 0.10 | 128 | 44 | 0.26 |

**Supplemental Table S3:** Beta regressions of mean proportion of flowers on a plant receiving pollen or hosting microbes by plant-level mean floral trait values. Plant-level data were calculated from open flower data (study 2).

| Proportion of flowers receiving Pollen | Fixed Effects | p |
| --- | --- | --- |
| Conspecific | Nectar Concentration | 0.091 |
| Conspecific | Nectar Volume | 0.63 |
| **Conspecific** | **Flower Length** | **0.011** |
| Conspecific | Flower Width | 0.12 |
| Heterospecific | Nectar Concentration | 0.78 |
| Heterospecific | Nectar Volume | 0.089 |
| Heterospecific | Flower Length | 0.71 |
| Heterospecific | Flower Width | 0.26 |
| Proportion of flowers containing Microbes | Fixed Effects | p |
| Bacteria | Nectar Concentration | 0.15 |
| **Bacteria** | **Nectar Volume** | **0.038** |
| Bacteria | Flower Length | 0.92 |
| Bacteria | Flower Width | 0.13 |
| Yeasts | Nectar Concentration | 0.28 |
| Yeasts | Nectar Volume | 0.75 |
| **Yeasts** | **Flower Length** | **< 0.001** |
| **Yeasts** | **Flower Width** | **< 0.001** |

**Supplemental Figure 1:**
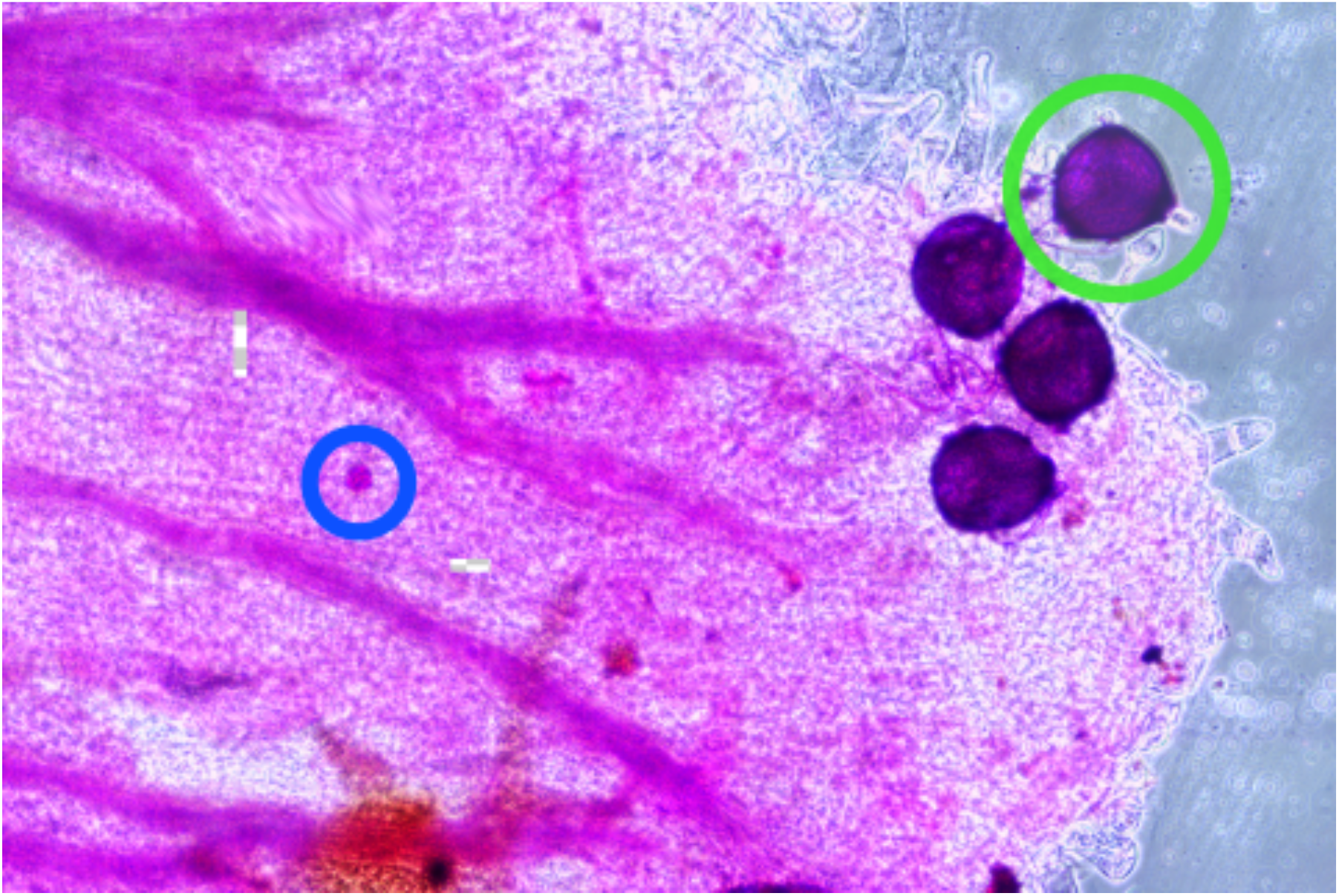
*Epilobium canum* stigma stained with basic fuchsin (digitally fused microscopic panorama at 20x). *Epilobium* pollen tetrad (4 pollen grains) circled in green, and heterospecific pollen grain circled in blue.

**Supplemental Figure 2:**
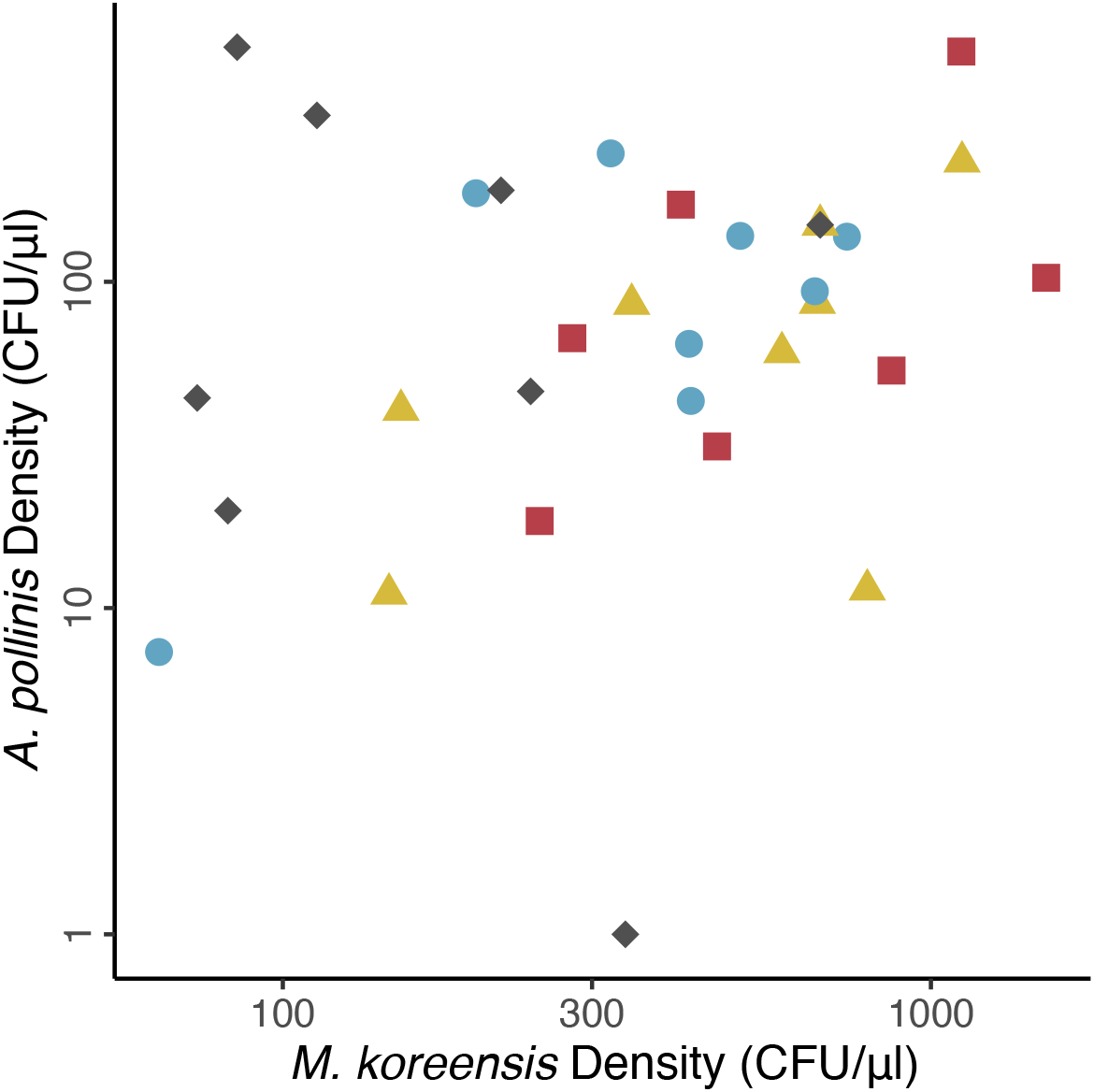
Scatterplot of mean final densities of *A pollinis* (x) and *M. koreensis* (y) for each individual plant. There was no correlation between the final densities of these microbes at the plant level. (Silver: grey diamonds, Canum: blue circles, Everett’s: yellow triangles, Calistoga-red squares).

